## Supplementary Information for "Nanofluidic chips for cryo-EM structure determination from picoliter sample volumes"

### Nanofluidic sample cells for cryo-EM imaging

#### Supplementary Tables

S1 Data collection and refinement statistics for collected datasets . . . 2
S2 Model refinement statistics for ApoFtn atomic model. . . . 2
S3 Data and 3D reconstruction parameters of reference datasets . . . 3

#### Supplementary Figures

S1 Fabrication schematic of cryoChips . . . 4
S2 Preparation of cryoChips for TEM imaging . . . 5
S3 Kymographs showing liquid entry and exit through the supply- and exit microchannels . . . 6
S4 Vitrification efficiency analysis . . . 7
S5 Simulation of plunge freezing of a simplified model of cryoChips in COMSOL 5.4 . . . 8
S6 Single-particle analysis workflow for *P. furiosus apoferritin* (ApoFtn) . . . 9
S7 Single-particle analysis workflow for tobacco mosaic virus (TMV). . . . 10
S8 Single-particle analysis workflow for *T. acidophilum* T20S proteasome (T20S) . . . 11
S9 Beam-induced motion of ApoFtn dataset in cryoChips . . . 12
S10 Tomograms of cryoChips with TMV and T20S . . . 13
S11 Surface electrostatic potential maps. . . . 14

#### Supplementary Movies

SM1 Filling process of a cryoChip through the microcantilever.
SM2 Zoom-in movie of (1) entire chip, (2) observation membrane, (3) nanochannel and (4) single ApoFtn particle.

**Table S1.** Data collection and refinement statistics for collected datasets

| | **ApoFtn**<br>**EMD-12901** | **TMV**<br>**EMD-12903** | **T20S**<br>**EMD-12915** |
|---|---|---|---|
| Specimen | 3 cryoChips filled with 3.4 mg/mL ApoFtn | 2 cryoChips filled with 1.1 mg/mL TMV | 1 cryoChip filled with 1.4 mg/mL T20S proteasome |
| Microscope | TFS Titan Krios | JEOL 3200-FSC | JEOL 3200-FSC |
| Voltage (kV) | 300 | 300 | 300 |
| Detector | K2-XP | K2-XP | K2-XP |
| Energy filter | Gatan BioQuantum | In-column omega filter | In-column omega filter |
| Number of movies (vitreous/crystalline) | 945 (815 / 130) | 82 (43 / 39) | 121 (91/30) |
| Pixel size (Å) | 0.8127 | 1.288 | 1.288 |
| Electron exposure ($e^{-}$/Å$^2$) | 63 | 59.7 | 59.7 |
| # of frames | 90 | 73 | 73 |
| Exposure time (s) | 9 | 10.95 | 10.95 |
| Defocus range (µm) | -1 to -2 | -1 to -3.5 | -2 to -4 |
| Symmetry imposed | O | helical rotation: 22.036°, helical rise: 1.415 Å | D7 |
| Map sharpening B-factor (Å$^2$) | -88.3 | -65.8 | -353.5 |
| Final number of particles / asymmetric units | 21,238 / 509,712 | 14,238 / 284,760 | 5750 / 80,500 |
| Final map resolution (Å) | 3.0 | 3.7 | 5.4 |
| Map resolution range (Å) | 2.9-3.5 | 3.5-4.3 | 5.2-6.5 |
| EMPIAR | 10708 | 10708 | 10708 |

**Table S2.** Model refinement statistics for ApoFtn atomic model.

| | **ApoFtn**<br>**7ohf** |
|---|---|
| Initial model used (PDB code) | 2x17 |
| Model resolution (Å, FSC = 0.5) | 3.3 |
| CC mask | 0.776 |
| **Model composition** | |
| Nonhydrogen atoms | 24 x 1399 |
| Protein residues | 24 × 169 |
| **RMSD** | |
| Bond lengths (Å) | 0.008 |
| Bond angles (°) | 0.864 |
| **Validation** | |
| MolProbity score | 0.88 |
| Clashscore | 1.41 |
| Rotamer outliers (%) | 1 |
| **Ramachandran plot** | |
| Favored (%) | 99.4 |
| Allowed (%) | 0.6 |
| Disallowed (%) | 0 |

**Table S3.** | **Data and 3D reconstruction parameters of reference datasets.**

| EMPIAR | Protein | Support film | Resolution | B-factor | # particles | Pixel size (Å) | Acq. scheme | Ref. |
|---|---|---|---|---|---|---|---|---|
| 10200 | ApoF | QF-R2/2 holey carbon | 1.65 Å | 66 $Å^2$ * | 426,450 | 0.814 | multi-shot | (44) |
| 10272 | ApoF | Ultrafoil-R0.6/1 + graphene | 2.14 Å | 54 $Å^2$ ** | 41,202 | 0.649 | single-shot | (39) |
| 10306 | TMV | CF-2/2-2C holey carbon | 1.90 Å | 41 $Å^2$ ** | 20,000 | 0.638 | multi-shot | (67) |
| 10389 | Urease | QF-R1.2/1.3 holey carbon | 1.98 Å | 38 $Å^2$ ** | 119,020 | 0.639 | both | (68) |
| 10708 | ApoFtn | cryoChip $SiN_x$ | 2.99 Å | 88.3 $Å^2$ ** | 21,238 | 0.813 | multi-shot | this paper |

*B-factor from ResLog plot. **B-factors as determined from Guinier plot.
ApoF: horse spleen apoferritin; ApoFtn: *P. furiosus* apoferritin

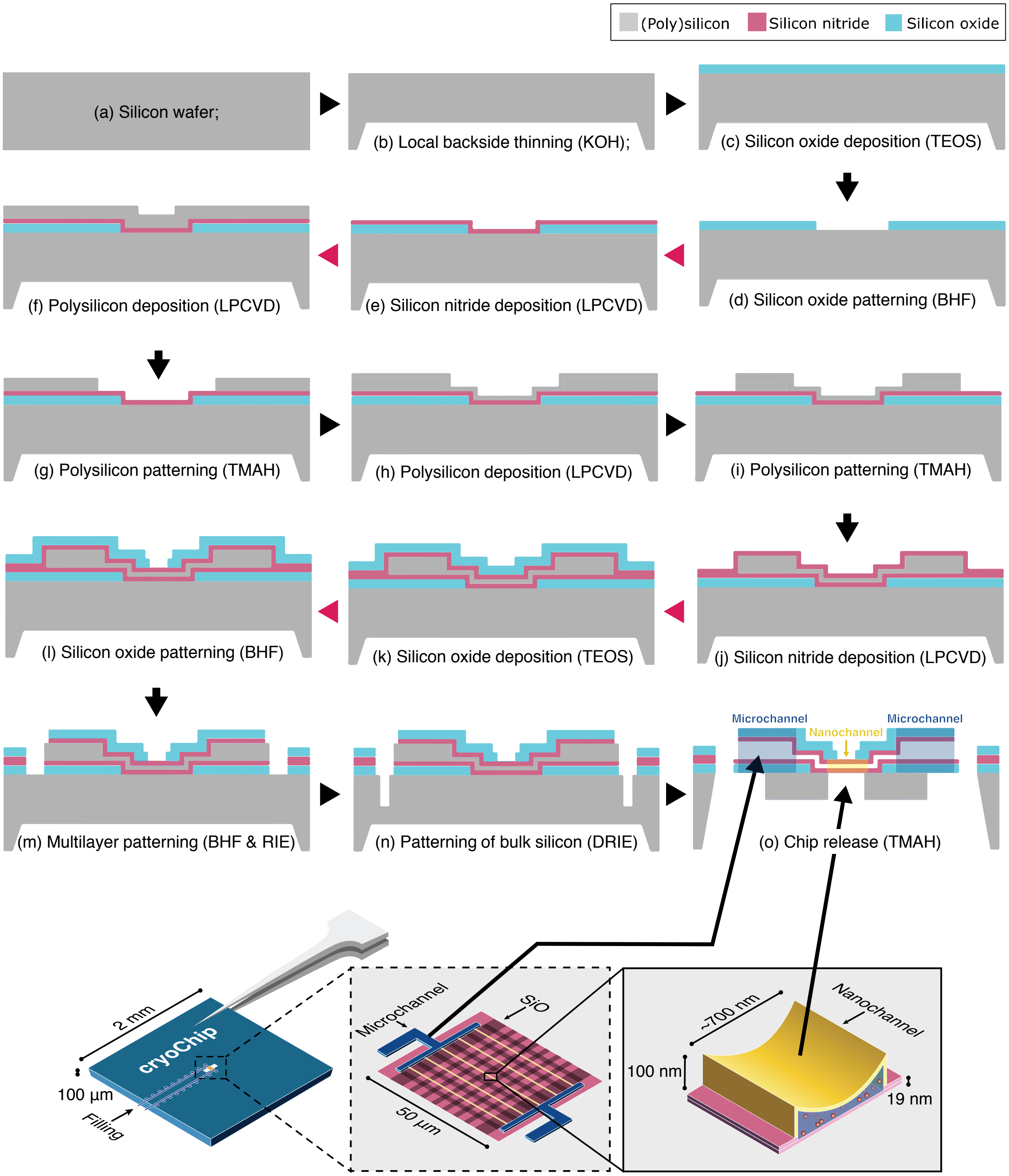


**Fig. S1.** | **Fabrication schematic of cryoChips.** All steps for nanofabrication of the cryoChip are shown in cross-section. The graphic on the bottom shows the location of microchannels (supply channels for filling) and nanochannels (for imaging) in the schematic.

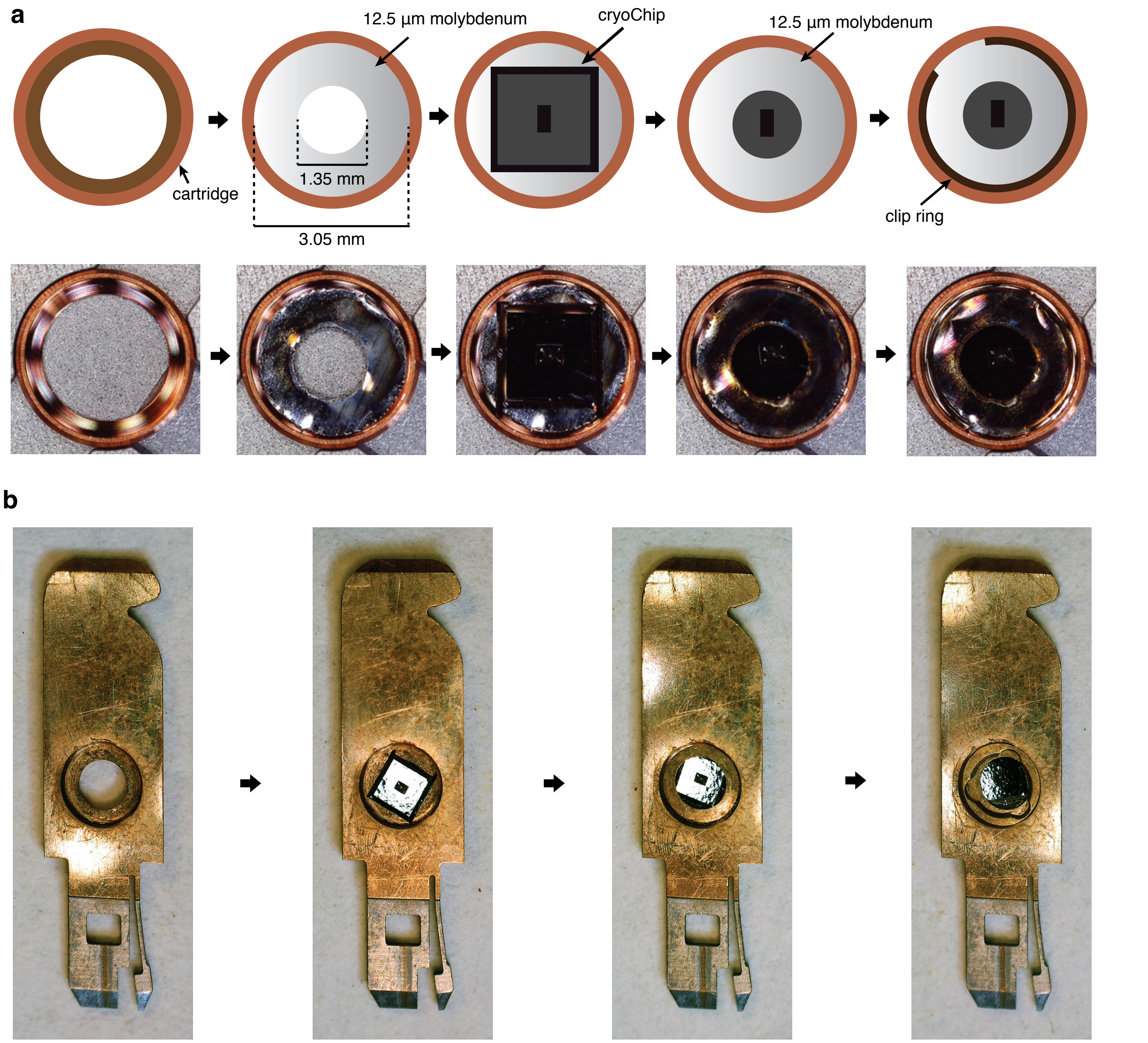


**Fig. S2.** | **Preparation of cryoChips for TEM imaging.** (a) Mounting in Autogrid cartridges (Thermo Fisher Scientific). A cryoChip is sandwiched between two molybdenum adapter rings with a circular aperture (laser cut from 12.5 μm molybdenum foil) and fixated with the Autogrid clip ring. Chips are mounted with the observation membrane facing the cartridge base and the slanted KOH-etch side walls facing the clip ring (b) Mounting in cartridges for the JEOL JEM3200-FSC microscope. The chip is placed with the observation membrane facing the cartridge base. The default metal spacer ring is placed on top and the assembly is secured by a screw ring.

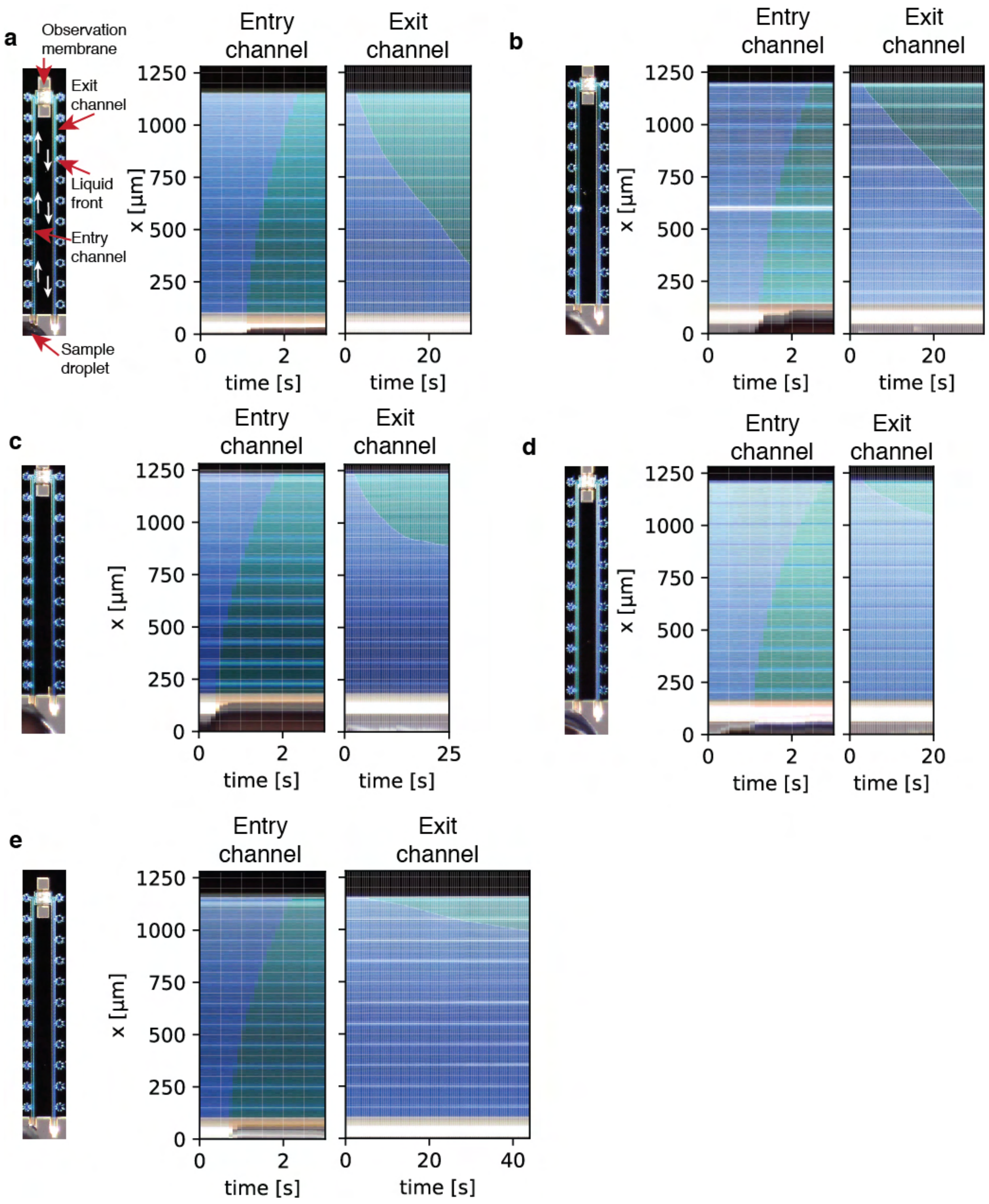


**Fig. S3.** | **Kymographs showing liquid entry and exit through the supply- and exit microchannels.** The liquid takes around one second to flow from the cantilever inside the whole entry channel. Due to the small cross-section of the nanochannels in the observation membrane, liquid flows out of the exit channel very slowly.

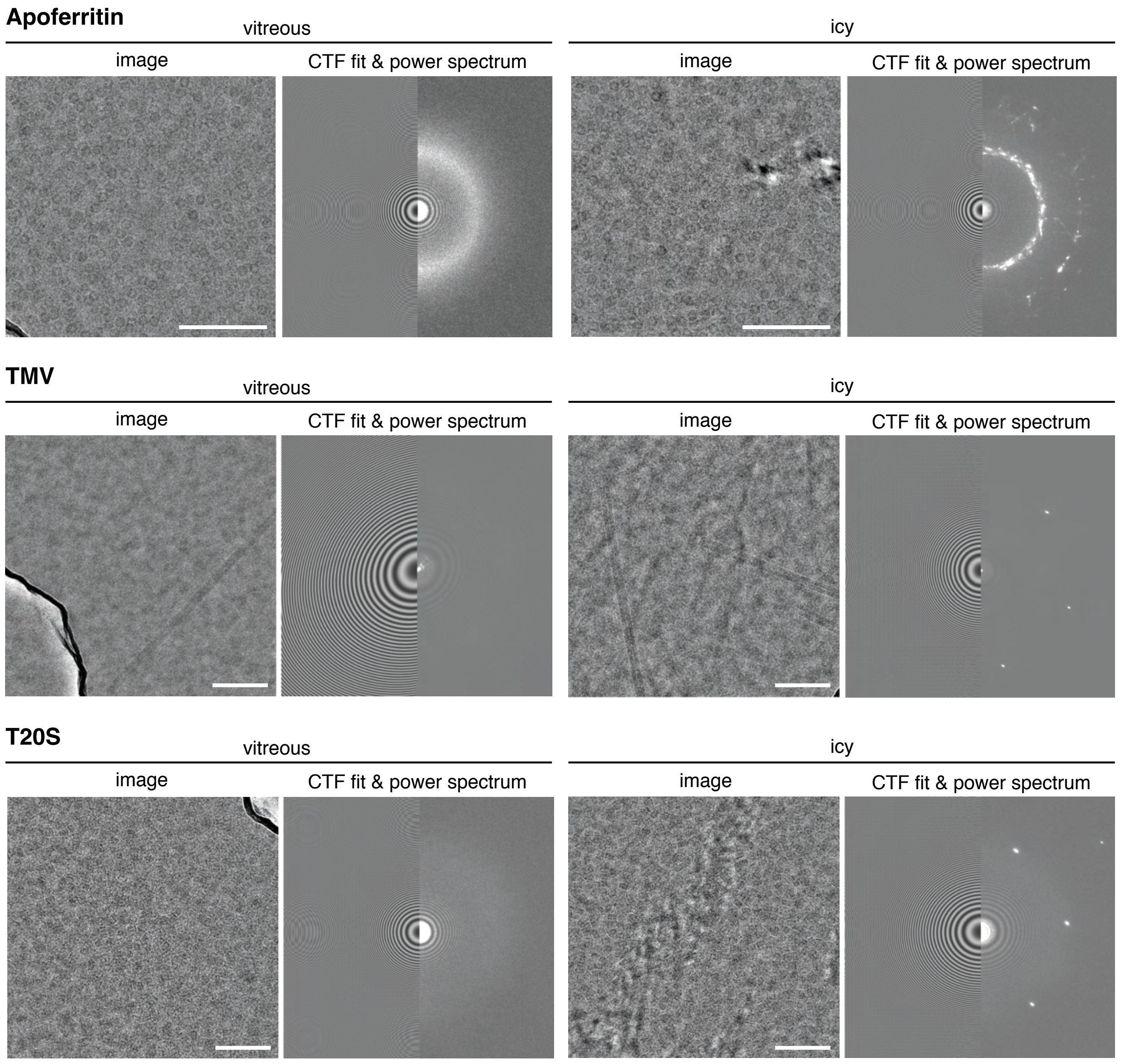


**Fig. S4.** | **Vitrification efficiency analysis.** Representative images with vitreous ice (left panels) and crystalline ice (right panels) from the three test specimens. CTF fits to the power spectra are also shown. Thon rings from $SiN_x$ can be used for coma-free alignment. Scale bar is 100 nm.

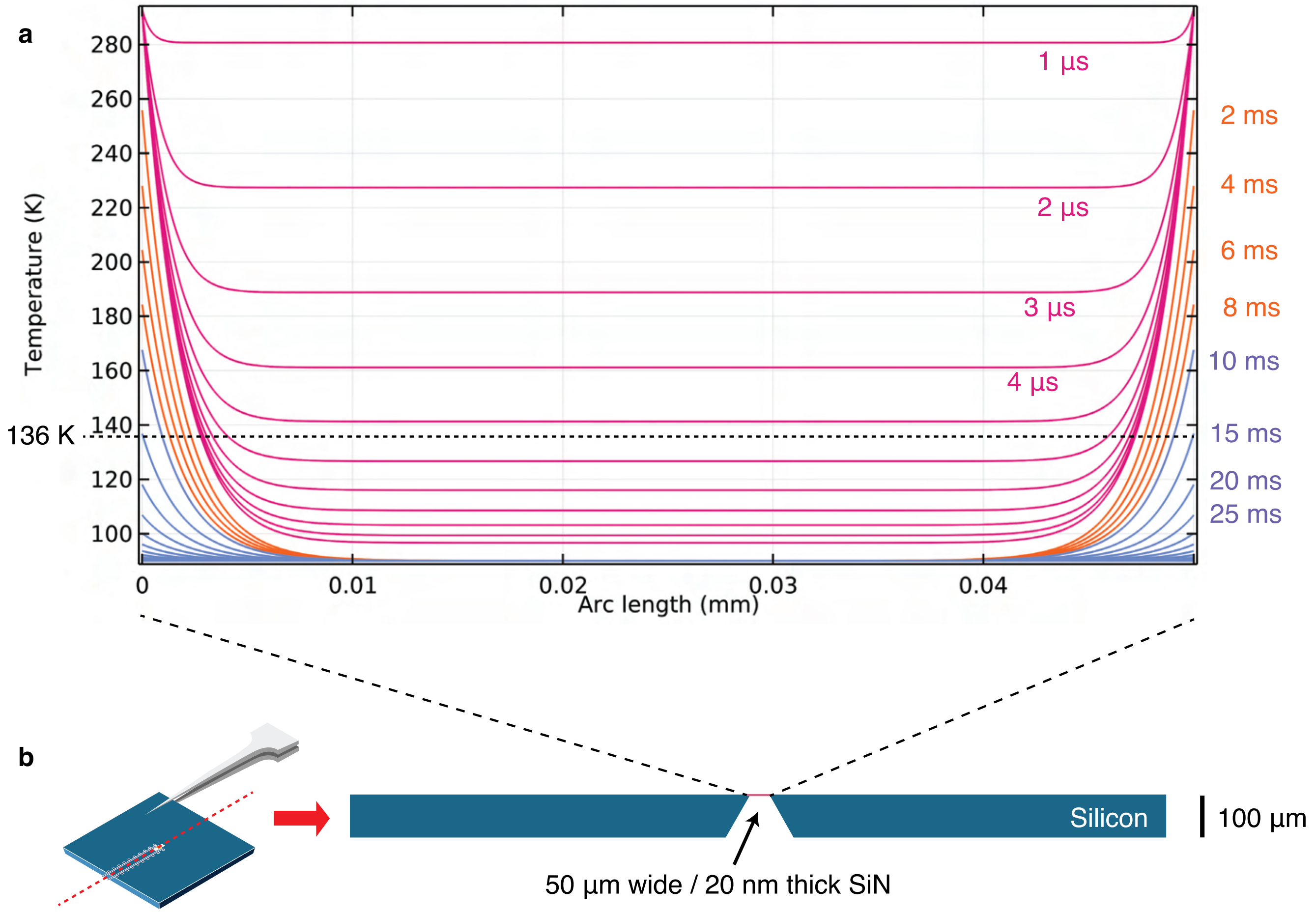


**Fig. S5.** | **Simulation of plunge freezing of a simplified model of cryoChips in COMSOL 5.4.** (a) Temperature profiles over 50 µm observation membrane from 1-10 µs (pink), 1-10 ms (orange) and 10-100 ms (blue). Dashed line shows glass-transition temperature of water at 136 K. (b) 2D model of cryoChips with 2 mm x 100 µm silicon base and 50 µm x 20 nm $SiN_x$ observation membrane used in simulations.

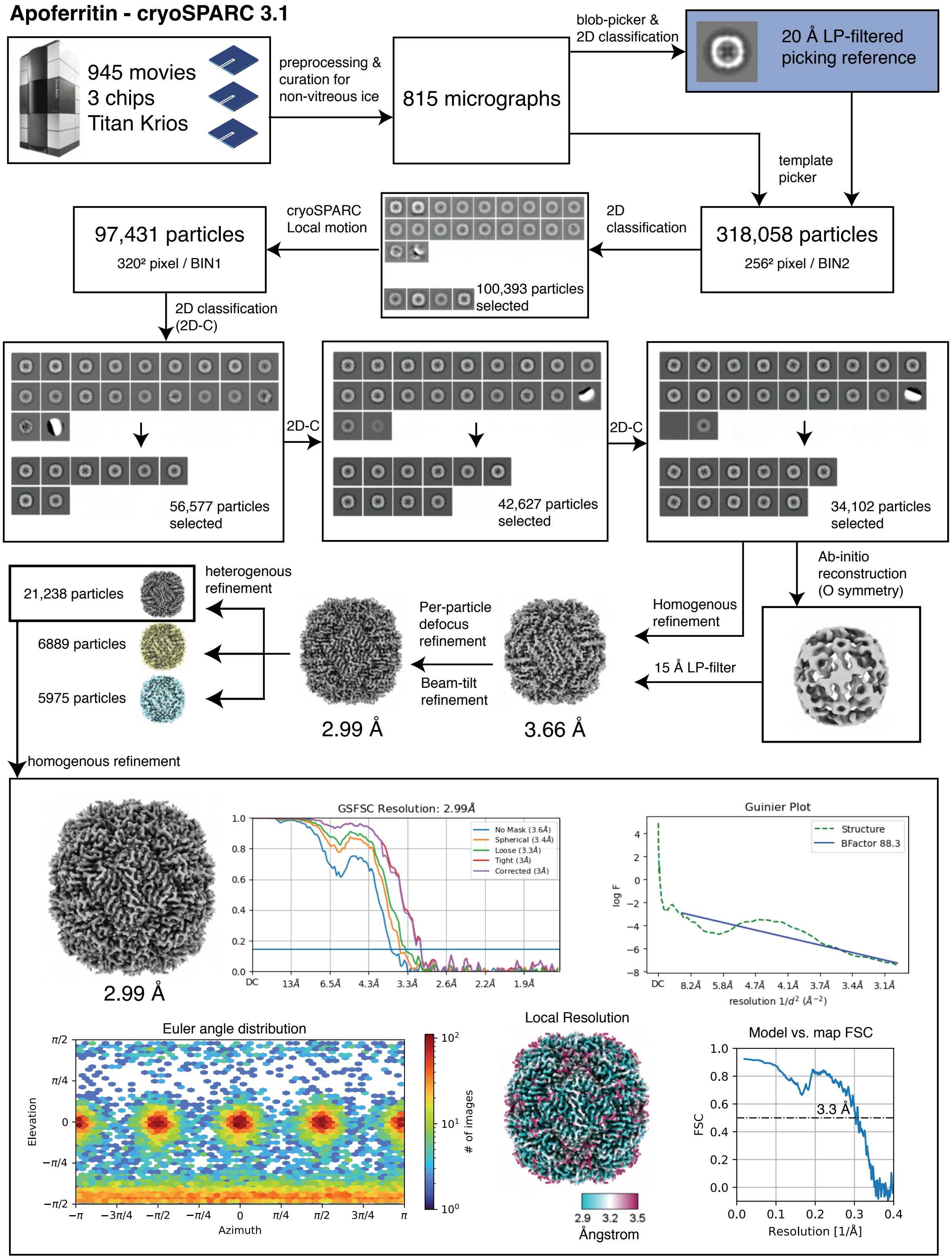


**Fig. S6.** | **Single-particle analysis workflow for *P. furiosus apoferritin* (ApoFtn).**

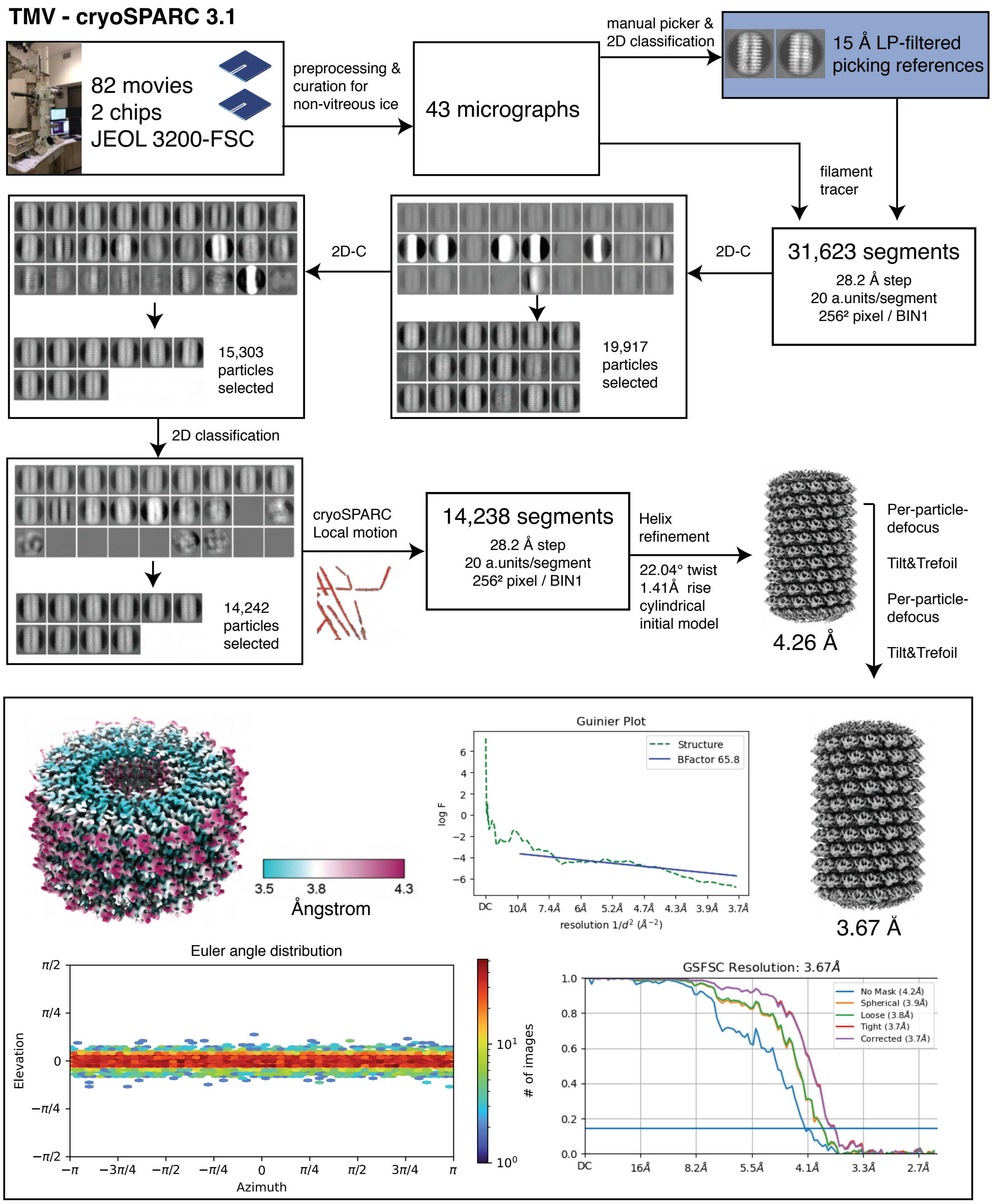


**Fig. S7.** | **Single-particle analysis workflow for tobacco mosaic virus (TMV)..**

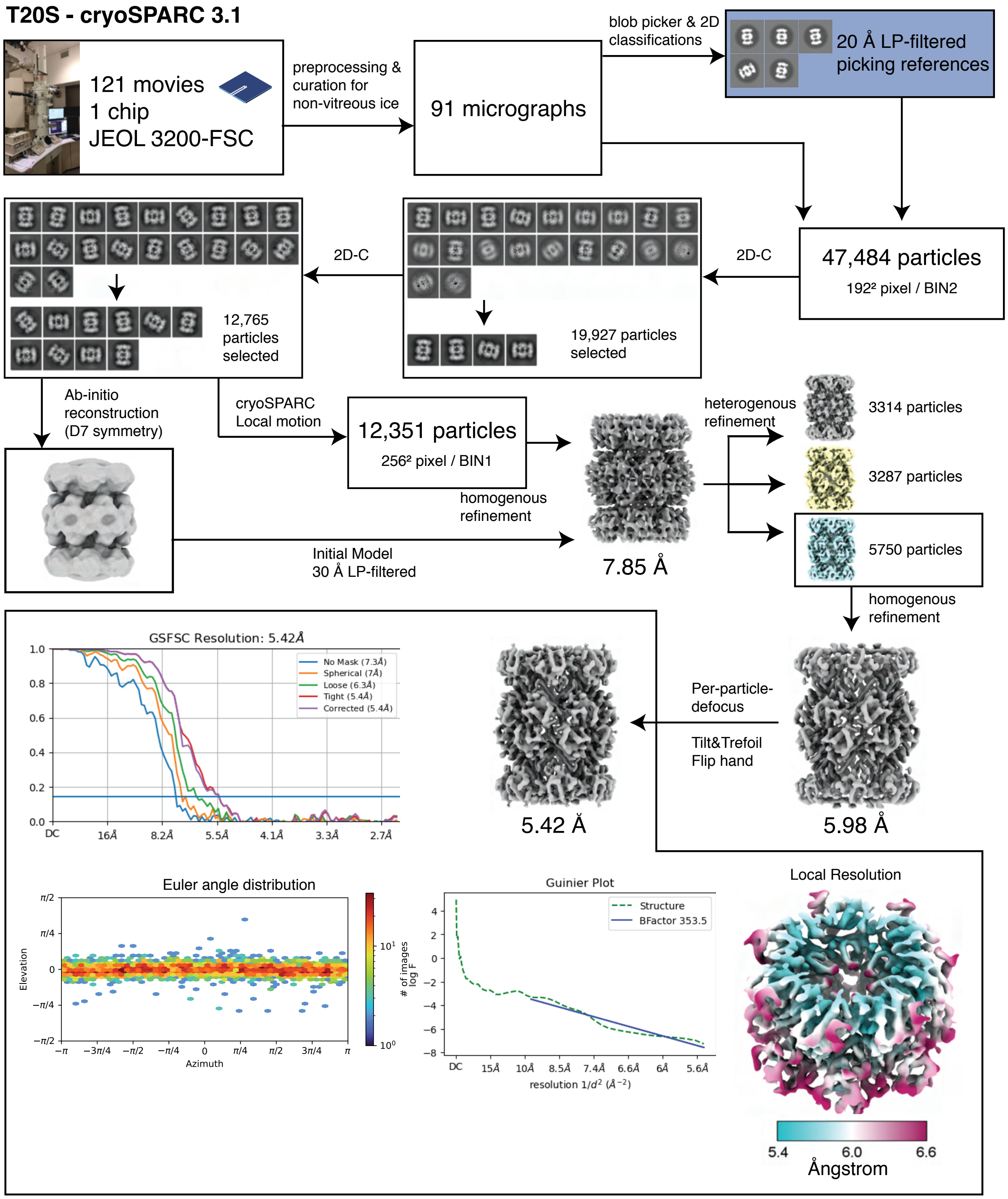


**Fig. S8.** | **Single-particle analysis workflow for *T. acidophilum* T20S proteasome (T20S).**

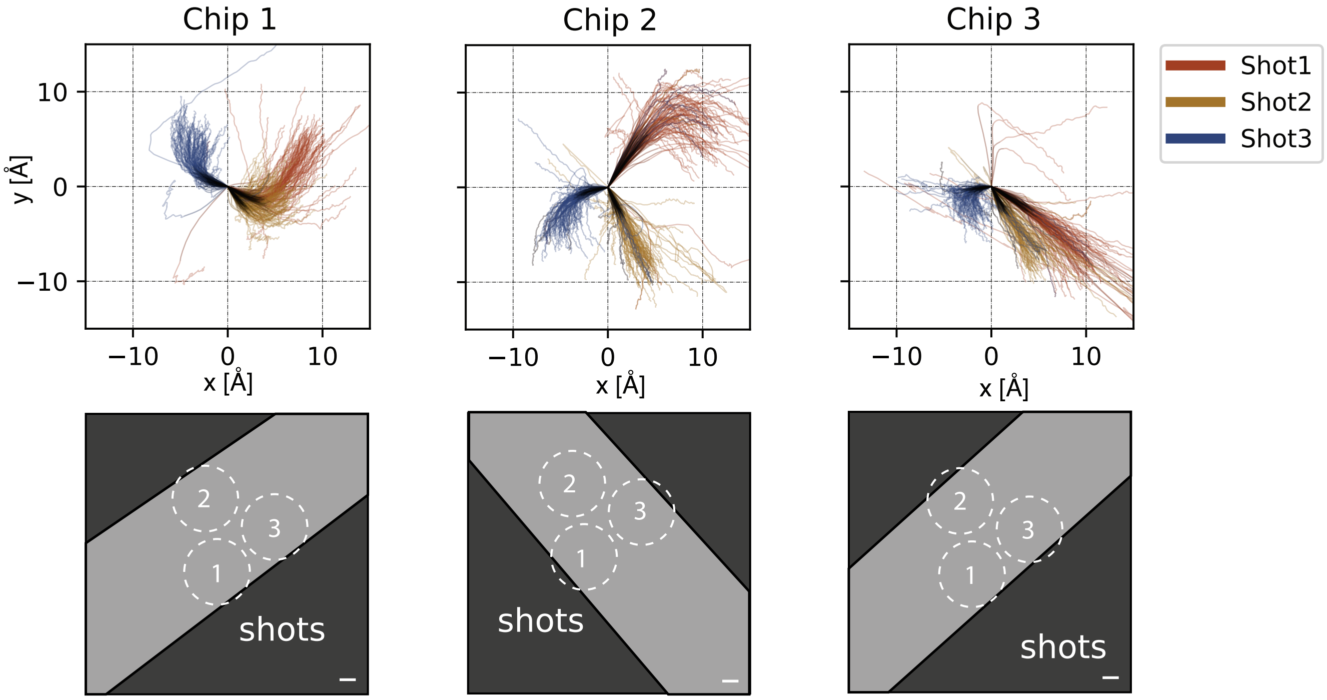


**Fig. S9.** | **Beam-induced motion of ApoFtn dataset in cryoChips.** Motion trajectories separated by chip and colour-coded by acquisition location reveal preferred directions of movement specific for each shot position in the nanochannels. Pictograms show the direction of nanochannels in the acquisition movies. Scale bar: 100 nm.

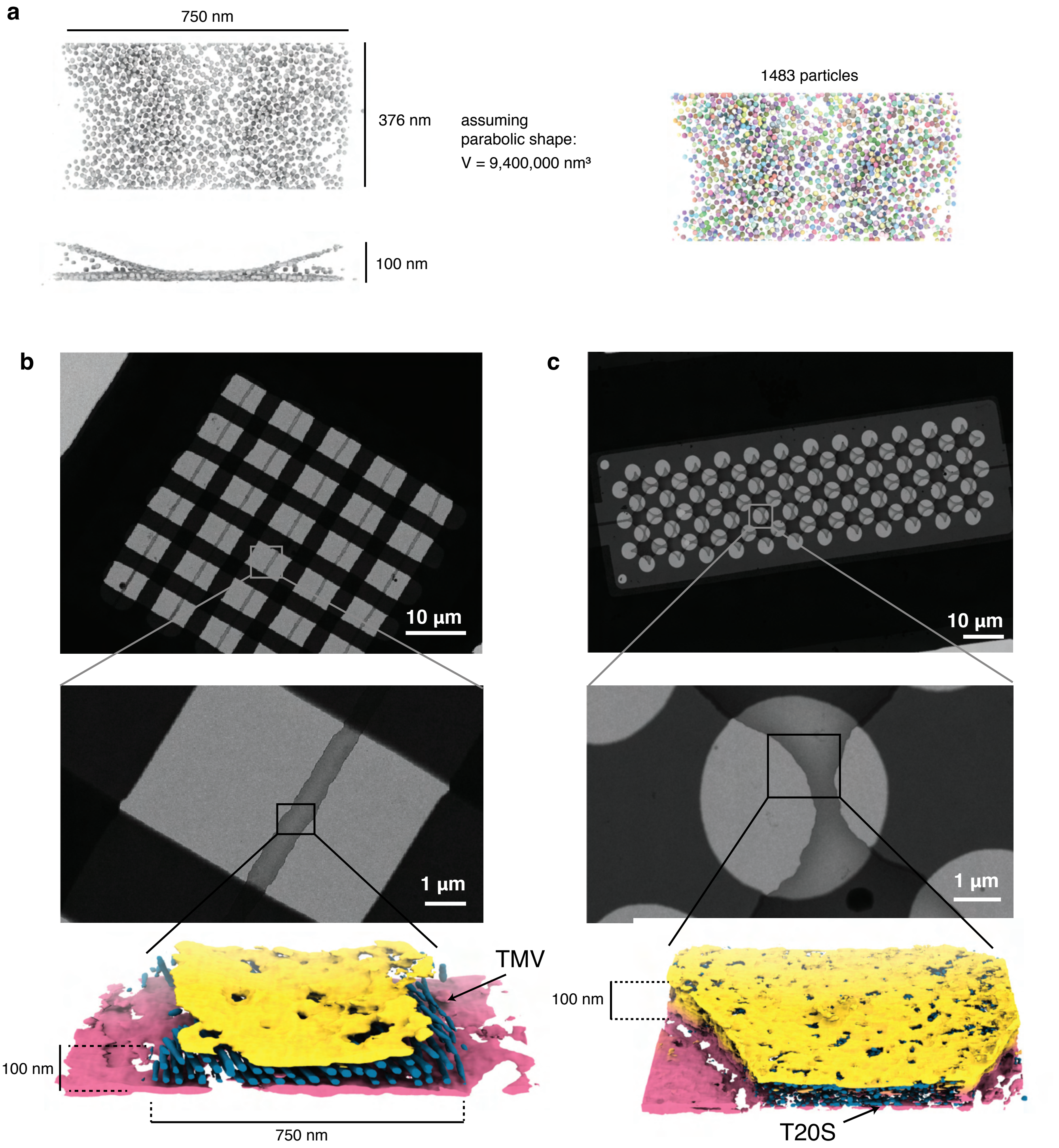


**Fig. S10.** | **Tomograms of cryoChips with TMV and T20S.** (a) Schematic illustration of volume determination and particle counting for a part of the ApoFtn tomogram. Segmented ApoFtn particles are shown as spheres. Low (top) and medium (center) magnification micrographs and segmented tomograms of nanochannels filled with (b) TMV (EMD-12914) and (c) T20S proteasome (EMD-12917). The cryoChip for T20S has a clover-patterned design of nanochannel and circular observation windows.

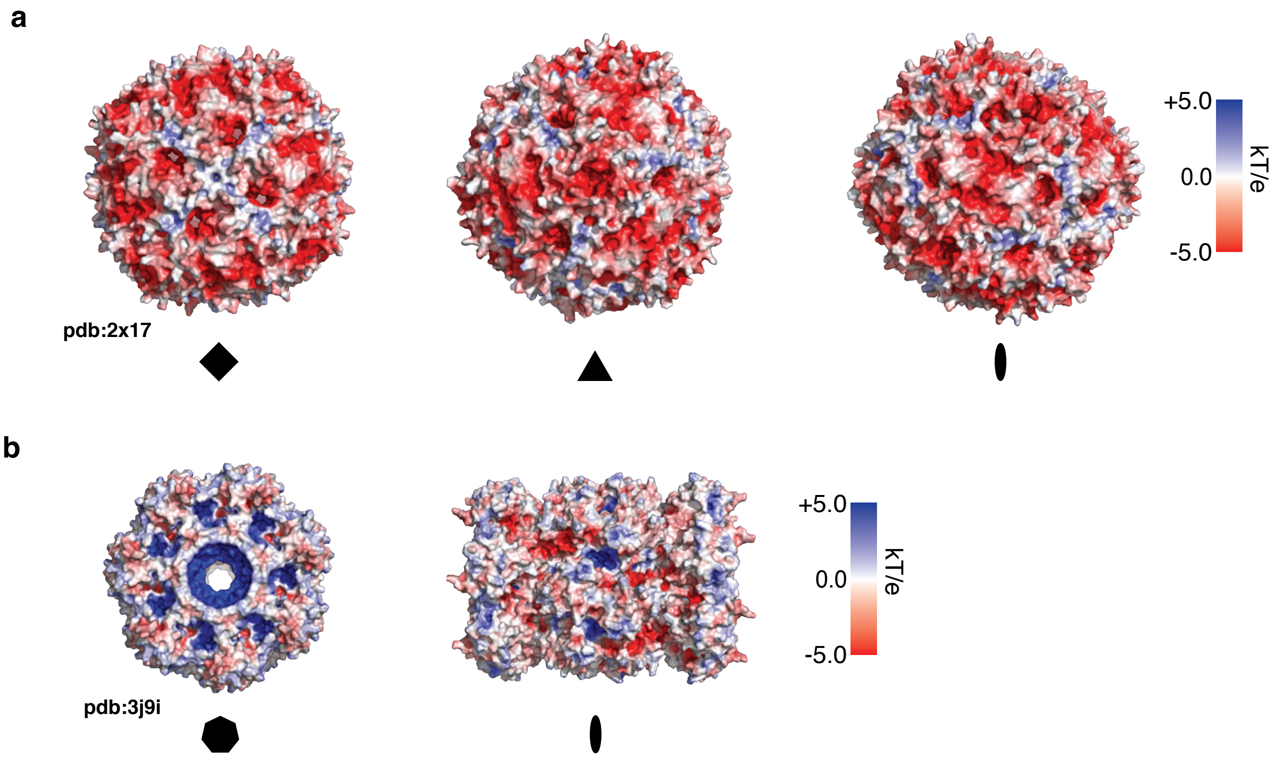


**Fig. S11.** | **Surface electrostatic potential maps..** Electrostatic potential maps for (a) ApoFtn, displayed along four-fold, three-fold and two-fold axes and (b) T20S proteasome displayed along the seven-fold and two-fold axes. Electrostatic potentials have been computed from PDB ID 2x17 (30) and PDB ID 3j9i (32) using Adaptive Poisson Boltzmann server (ABPS) (71).
